# Improving Protein Structure Prediction Using Integrative Cryo-EM and Ion Mobility Mass Spectrometry Modeling

**DOI:** 10.64898/2026.02.07.704481

**Authors:** Jacob B. Howard, Akshaya Narayanasamy, Steffen Lindert

**Affiliations:** Department of Chemistry and Biochemistry, Ohio State University, Columbus, OH, USA; Department of Chemistry and Biochemistry, University of California, Los Angeles, Los Angeles, CA, USA

## Abstract

Proteins perform essential roles across nearly all cellular processes, and accurate three-dimensional structures remain critical for elucidating structure–function relationships and studies on drug discovery. Cryo-electron microscopy (cryo-EM), X-ray crystallography, and nuclear magnetic resonance can provide detailed structural information. However, for many proteins, structural information is available only as lower-resolution experimental data or sparse data. Such information is more difficult to translate into accurate atomic coordinates; a common example is low-resolution cryo-EM density maps. In parallel, mass spectrometry-based methods, including ion mobility (IM-MS), offer rapid, broadly applicable structural descriptors such as collisional cross section (CCS), a global measure of molecular shape and size, but CCS values also do not provide atomistic detail. Here we present CRIM (cryo-EM + IM-MS), an integrative Rosetta scoring function that combines low-resolution cryo-EM density information with IM-MS derived CCS as restraints to improve monomeric protein structure prediction. CRIM incorporates the Rosetta REF2015 (RS) energy with a CCS agreement penalty (computed via PARCS) and an electron-density agreement term (elec_dens_fast). We tested CRIM on an ideal dataset of 60 monomeric proteins using simulated CCS values and density maps. Across the ideal dataset, the CRIM score function improved or maintained prediction quality for many targets, reducing the mean RMSD from 3.65 Å (RS) to 2.90 Å and increasing the mean TM-score from 0.88 to 0.90. Furthermore, an experimental benchmark dataset of 54 proteins was curated to include either experimental cryo-EM maps or published CCS values. On the experimental dataset, CRIM similarly improved model selection, lowering the mean RMSD from 6.65 Å to 4.38 Å and raising the mean TM-score from 0.73 to 0.79. In comparison to AlphaFold3 predictions, CRIM frequently yielded competitive predictions and was able to substantially outperform AlphaFold3 for select difficult targets where sparse experimental restraints provide strong discriminatory power. The CRIM score function is freely available within the Rosetta software suite and provides a practical framework for leveraging complementary IM-MS and cryo-EM data to improve monomeric protein structure prediction.

## Introduction

Proteins are fundamental for effectively all cellular processes, serving as enzymes, structural components, and signaling molecules. Understanding their three-dimensional structure is crucial for elucidating their functions and applications in drug discovery^1, 2^, vaccine design^3^, and other biotechnological applications^4^. High-resolution techniques such as X-ray crystallography, nuclear magnetic resonance (NMR) spectroscopy, and cryo-electron microscopy (cryo-EM) are instrumental in determining protein structures at atomic resolutions^5^. While effective, these methods often require extensive sample preparation, large quantities of material, and may not be suitable for all proteins when solving structures at high resolutions. Occasionally, some of these higher resolution techniques only yield structural information at lower resolutions, such as when lower-resolution density maps are obtained from cryo-EM, or when NMR experiments yield sparse data^6-10^.

In addition to high-resolution approaches, a variety of experimental techniques provide sparse yet valuable structural information. Mass spectrometry (MS) is a prominent example, offering rapid data collection, minimal sample requirements, and broad applicability across a wide range of protein sizes and heterogeneous samples. Several MS-based methods are routinely employed to probe structural features, including hydrogen/deuterium exchange (HDX), chemical cross-linking (XL-MS)^11^, surface-induced dissociation (SID)^12^, covalent labeling (CL), and ion mobility (IM-MS)^13, 14^. These approaches yield complementary insights into protein architecture, such as connectivity, topology, stoichiometry, post-translational modifications, and ligand binding. IM-MS provides a collisional cross section (CCS) value, which reflects the overall size and shape of an ion based on a rotationally averaged cross-sectional area, thereby enabling characterization of conformational states. Although the structural information from these lower-resolution methods is insufficient to resolve atomic coordinates, it nonetheless offers critical constraints that may complement high-resolution or computational techniques.

As an alternative method, computational techniques can be used to predict protein structure based solely on the amino acid sequence without any experimental data. Recent advances in machine learning, particularly with deep learning models like AlphaFold3 (AF3)^15^, Boltz2^16^, and RosettaFold2^17^, have significantly enhanced our ability to predict protein structures. While these models are highly effective and frequently yield accurate predictions, there is a significant number of proteins that remain challenging to predict^18^. To improve computational structure prediction, integrating sparse experimental data into the prediction process can further improve the accuracy of these predictions^19^. In these cases, experimental data serves as constraints in the model, helping to direct and refine predictions.

Previous studies have demonstrated that diverse experimental data types can be incorporated into modeling frameworks such as Rosetta to improve protein structure predictions^20-22^. For example, XL-MS data have been used as upper-bound distance restraints that constrain the relative positioning of residues within a protein complex^23, 24^. CL experiments contribute information on residue solvent accessibility, providing complementary constraints that refine surface exposure and topology in modeled structures^25-29^. SID measurements capture subunit connectivity and non-covalent interaction energetics, offering additional constraints on the stability and architecture of protein assemblies^30-34^. Similarly, IM-MS experiments provide collisional cross section (CCS) values that serve as a global shape descriptor, enabling discrimination between candidate models based on their overall shape and size^14, 35-41^. Beyond mass spectrometry, low-resolution cryo-EM density maps can be directly integrated into modeling protocols, supplying spatial localization constraints that further refine structural predictions^9, 10, 23, 32, 42-44^. Furthermore, integrative approaches that combine multiple complementary experimental techniques have been shown to be successful. Seffernick et al. demonstrated that integrating SID mass spectrometry data with cryo-EM maps improved the accuracy of protein complex modeling beyond what was observed when just SID or cryo-EM was used independently^32^. Similarly, studies combining ion mobility with chemical cross-linking have successfully refined modeling of large protein assemblies by simultaneously constraining both overall shape and specific residue-residue distances^45^. These examples illustrate how incorporating diverse experimental datasets into computational modeling frameworks can yield more robust and biologically meaningful structural predictions^46^.

In this study, we extend these integrative approaches to the problem of monomeric protein structure prediction. Building on the evidence that combining complementary data types further improves structural models, we hypothesized that combining sparse IM-MS and cryo-EM data with RosettaCM modeling could help improve the structure prediction of monomeric proteins. These two sources of information are complementary, as CCS values from IM-MS provide a global description of molecular size and shape, whereas cryo-EM density maps capture three-dimensional spatial localization. CCS measurements are rotationally averaged, in contrast to orientation-specific information from cryo-EM, which allows the two methods to constrain models in distinct and orthogonal ways. To leverage this complementarity, we present the CRIM (cryo-EM + IM-MS) score function which incorporates sparse data obtained from low-resolution (>6 Å) cryo-EM density map and IM-MS in the form of CCS values into Rosetta comparative modeling (CM) to improve protein structure predictions. With our method we were able to improve structure predictions across 83% of the monomeric proteins in both our datasets from an average RMSD of 5.1 Å with the Rosetta REF2015 (RS) score function to 3.6 Å RMSD with the new CRIM score function. The score function is part of the Rosetta software suite and is freely available for access by the users. (GitHub link: https://github.com/RosettaCommons/rosetta/).

## Methods

### Ideal Dataset Construction

To evaluate the performance of the CRIM scoring function for idealized, noise- and error-free data, we used a previously assembled ideal dataset of 60 monomeric proteins^40^. Structures in the ideal dataset represent all unique protein architecture categories as classified by the CATH Protein Structure Classification Database, with sequence lengths ranging from 58 to 965 amino acids. For each protein, synthetic experimental data was generated: collision cross section (CCS) values were calculated using the Rosetta PARCS algorithm^40, 47^ and low-resolution cryo-electron microscopy (cryo-EM) density maps were simulated using the BCL density module as described below.

Structural models for each of the 60 proteins in ideal dataset were generated using Rosetta multi-template Comparative Modeling (CM)^48^. Following established protocols, 10,000 models were produced for each protein^40^. The 3mer and 9mer fragment libraries were created using the Rosetta fragment picker tool^49^. BLAST was used to identify both low and high sequence similarity templates which were included to broaden the diversity of the model set. Additionally, template weights were varied to produce both native-like (<5 Å RMSD) and non-native-like (>5 Å RMSD) models compared to the target protein. All structures generated with RosettaCM were subject to the Rosetta fast relax protocol prior to scoring.

Models were scored using the RS score function ^50^ and the CRIM function, respectively, to assess their ability to distinguish native-like and non-native-like conformations. Model quality was evaluated using alpha-carbon root-mean-square deviation (Cα-RMSD) calculated in PyMOL, and TM-score values were computed with TM-align^51^.

### Benchmark Dataset Construction

To evaluate CRIM under relevant experimental conditions, we curated a benchmark dataset of 54 monomeric proteins. Of these, 30 of the target proteins had experimentally determined cryo-EM density maps and 24 had published IM-MS CCS measurements, but no target had both data types. To enable integrated scoring across the full benchmark set, the missing data for each target was simulated: density maps were generated with BCL for targets with CCS data only, and CCS values were computed with PARCS for targets with cryo-EM data only. For each benchmark protein, 10,000 models were generated using the same RosettaCM protocol applied to the ideal dataset.

### Low-Resolution Cryo-EM Density Map Preparation

Within the benchmark dataset, 30 proteins had experimental cryo-EM density maps. However, the density maps were often for entire complexes, which would affect the scoring process of monomeric proteins when using the “elec_dens_fast” score term. As a solution, the density maps were cropped within 5 Å of the native structure using ChimeraX^52^.

For the remaining targets in the benchmark dataset, along with all the proteins in the Ideal dataset, the density maps were simulated using the BCL density package. All density maps were simulated from crystal structures using BCL::density::FromPDB^53^ at 14 Å resolution, with a voxel size of 1.4 Å and a Gaussian noise level set to 0.8.

### Native CCS Calculations for Proteins Without IM-MS Data

Theoretical CCS values were computed from atomic coordinates using the Rosetta PARCS algorithm^40^. Each structure was randomly rotated 300 times to obtain a rotationally averaged CCS. CCS calculations were performed with the buffer-gas radius set to 1.0 Å to approximate helium-based ion mobility conditions.

CCS values had to be calculated from the native structure for all 60 proteins in the ideal dataset and 30 proteins in the benchmark dataset. Several of these native proteins had missing residues within their PDB structure. Due to PARCS calculating CCS values based on the atoms present in a PDB file, missing residues in the structure may result in an underestimation of the native CCS values. For native structures with missing residues in their PDB entries, a modified RosettaCM pipeline was employed to reconstruct the full sequence across both datasets. For this RosettaCM protocol, only templates with high sequence similarity were utilized in addition to the crystal structure with the missing residues. Furthermore, the crystal structure was weighted higher than the other templates so that near-native-like predictions could be obtained. With this protocol, only 250 structures were predicted per protein system. The best-scoring model generated from this pipeline for each target protein was used to compute the theoretical CCS value using PARCS. We used this CCS value as the benchmark against the models in place of experimental IM-MS data.

### Protein Structure Prediction with AlphaFold3

For each protein in the ideal and benchmark datasets, structures were generated using AlphaFold3 v3.0.1 on the Cardinal cluster at the Ohio Supercomputer Center. Predictions were made using the standard AF3 settings. The highest-ranked AF3 model for each protein was used in comparison to the best-scoring CRIM predicted models

### CRIM Score Function

Density maps and CCS values from cryo-EM and IM-MS experiments, respectively, were incorporated into the RS score function to act as spatial restraints for integrative Rosetta modelling. The Cryo-EM and IM score terms provide information regarding a protein’s overall shape, size, and spatial density. To integrate this information into Rosetta, the IM_Score Term_ was adapted from a previously implemented score term to quantify the agreement of a predicted model with the IM-MS data^40^. The IM_Score Term_ is a piecewise penalty function (defined in Eq. 1) based on the difference (ΔCCS) between CCS_PARCS_ and CCS_IM_. The function uses a lower bound (LB) to account for error and an upper bound (UB) to cap the penalty for large deviations (shown in Eq. 1). ΔCCS values below the LB (25 Å^2^) do not receive a penalty and ΔCCS above the UB (125 Å^2^) receive the maximum penalty of 125, with a fade function used in between.

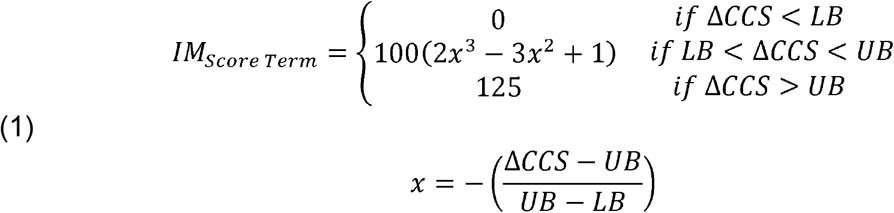

The cryo-EM_Score Term_ is a previously implemented score term in Rosetta (elec_dens_fast) and was calculated based on model agreement with either experimental cryo-EM density maps or the simulated density maps^54^.The evaluation score, CRIM_Score_, was defined as a weighted sum of the RS score function, IM_Score Term_ and cryo-EM_Score Term_ as shown in Eq. 2.

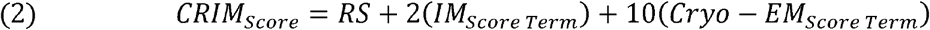

The total CRIM score captures both structural conformity to the standard Rosetta score, and quantitative agreement with sparse experimental data from cryo-EM and IM-MS. This integrated scoring strategy was hypothesized to improve the selection of near-native structures across datasets with diverse experimental resolutions and data completeness.

## Results and Discussion

### CRIM Score Function Improved Model Selection in an Ideal Dataset

In this study, we hypothesized that integrating cryo-EM density information with IM-MS data would provide complementary constraints. This integration would enable more accurate monomeric protein structure prediction than can been achieved with either method alone. As both cryo-EM and IM-MS are becoming more accessible and widely used, it is increasingly likely in the future that both data types will be available for the same protein target, underscoring the need for computational methods capable of leveraging these datasets together. To test this hypothesis, we assessed the performance of the CRIM score function using an ideal dataset of 60 monomeric proteins, designed to represent diverse protein folds across the CATH classification. In this dataset, synthetic experimental data was generated by simulating CCS values with the Rosetta PARCS algorithm^40^ and cryo-EM density maps with the BCL density package^10, 53^. For each protein, 10,000 model structures were generated with Rosetta comparative modeling (RosettaCM), and all models were subsequently rescored using the CRIM function to evaluate its ability to discriminate between native-like and non-native conformations.

For both scoring functions, the lowest-scoring model was considered the best or predicted structure. Upon incorporating cryo-EM and IM-MS data through the CRIM score function, we found a notable improvement in the accuracy of the best-scoring models compared to predictions made using only the RS score function. The structures predicted by CRIM exhibited an average RMSD of 2.90 Å, compared to 3.65 Å with the RS score function. The average TM-scores also improved, increasing from 0.88 with RS to 0.90 with CRIM. Three proteins demonstrating significant improvement in RMSD with the CRIM score function are illustrated in Figure 1. Figures 1A and 1B show that incorporating experimental data not only resulted in more accurate structural predictions but also penalized inaccurate conformations more strongly, yielding significantly more clearly defined energy funnels across the proteins. As illustrated in Figure 1C, the CRIM score function guided model selection toward native-like conformations for all three proteins. For example, hemopexin (PDBID: 1QHU) improved from 10.77 Å RMSD with RS to 3.78 Å with CRIM, while pORF2 (PDBID: 2ZZQ) improved from 11.75 Å to 4.46 Å. Even for the 50S ribosomal protein L15 (PDBID: 3CPW), which was one of the most challenging proteins in the ideal dataset, the CRIM score function reduced the RMSD from 22.61 Å to 5.44 Å, indicating that substantial gains can be achieved across diverse structural contexts. Similar results were observed for the other proteins in the ideal dataset as shown in Supplementary Fig. 1-3.

**Figure 1.**
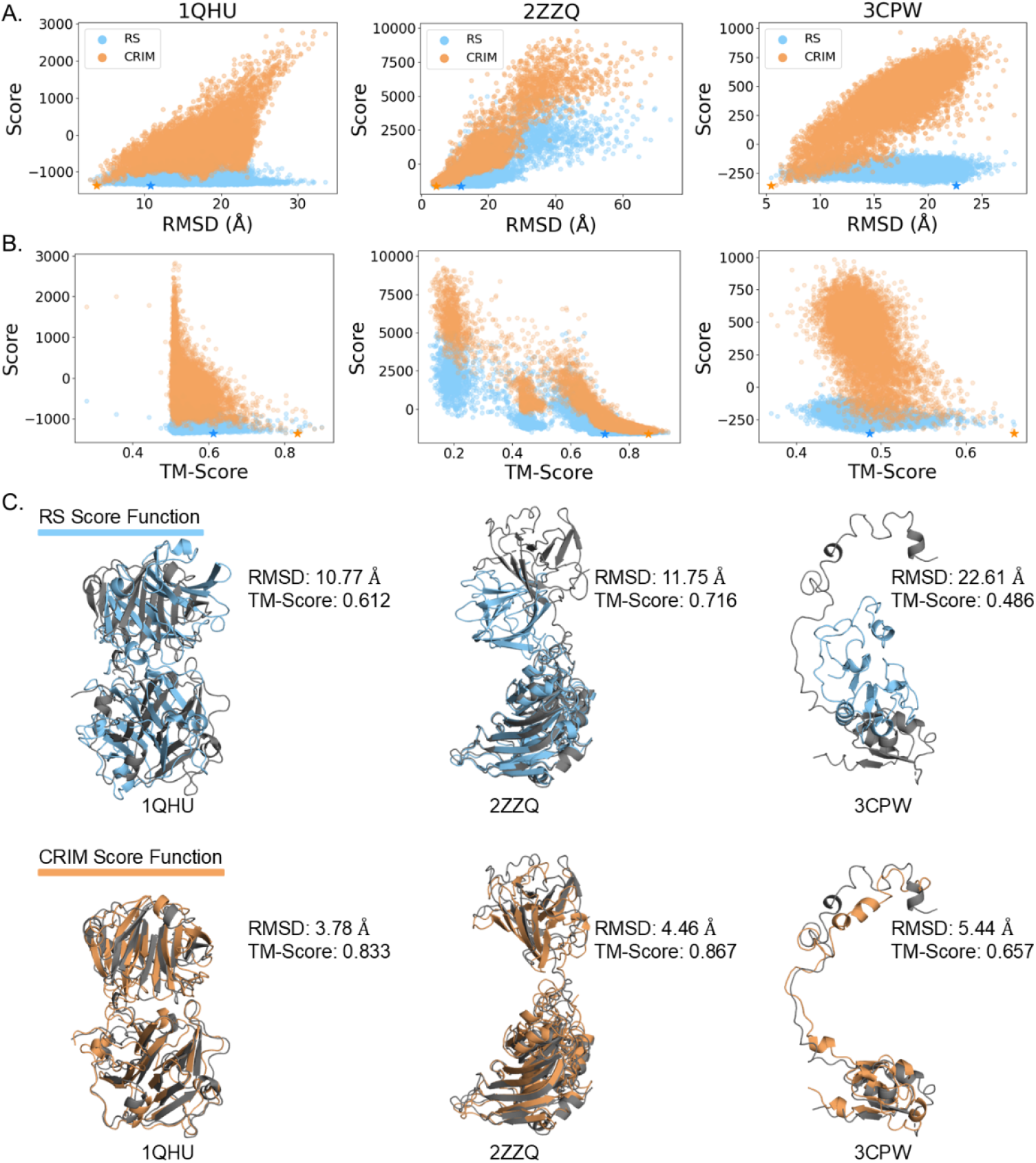
(A) The score vs RMSD distributions for the 10,000 model structures of proteins 1QHU, 2ZZQ, and 3CPW when scored with the CRIM score function (orange) compared to the RS score function (light blue). For both score functions, the best scoring structure is indicated with a star in its respective color. The score values for the CRIM score function were normalized to the RS scores by subtracting the difference between the two best scoring structures from all 10,000 of the model scores. (B) The corresponding score vs TM-score distributions of the models scored with the CRIM score function (orange) compared to the RS score function (light blue). (C) Predicted models for the three proteins using the RS score function (light blue) versus the CRIM score function (orange), each overlaid with the native structures (grey).

Across the ideal dataset, the RMSD of the predicted structures either improved or remained the same for 50 out of the 60 proteins, with an average improvement of 1.44 Å among the proteins that saw an improvement (Figure 2A (i)). Likewise, the TM-score improved for 49 out of 60 proteins with an average improvement of 0.04 over the RS score function for those proteins (Figure 2B (i)). Visually, most points lie below (RMSD) or above (TM-score) the x = y line, suggesting an improved prediction, and the deviation from the line grows at higher RS RMSD values, indicating stronger benefits for harder targets. These results further support that the addition of simulated cryo-EM and IM-MS data refines the ability of the RS score function to distinguish between native and non-native like models. However, since the simulated, idealized data is not identical to real experimental data, we further tested the effectiveness of the CRIM score function in conjunction with experimental data.

**Figure 2.**
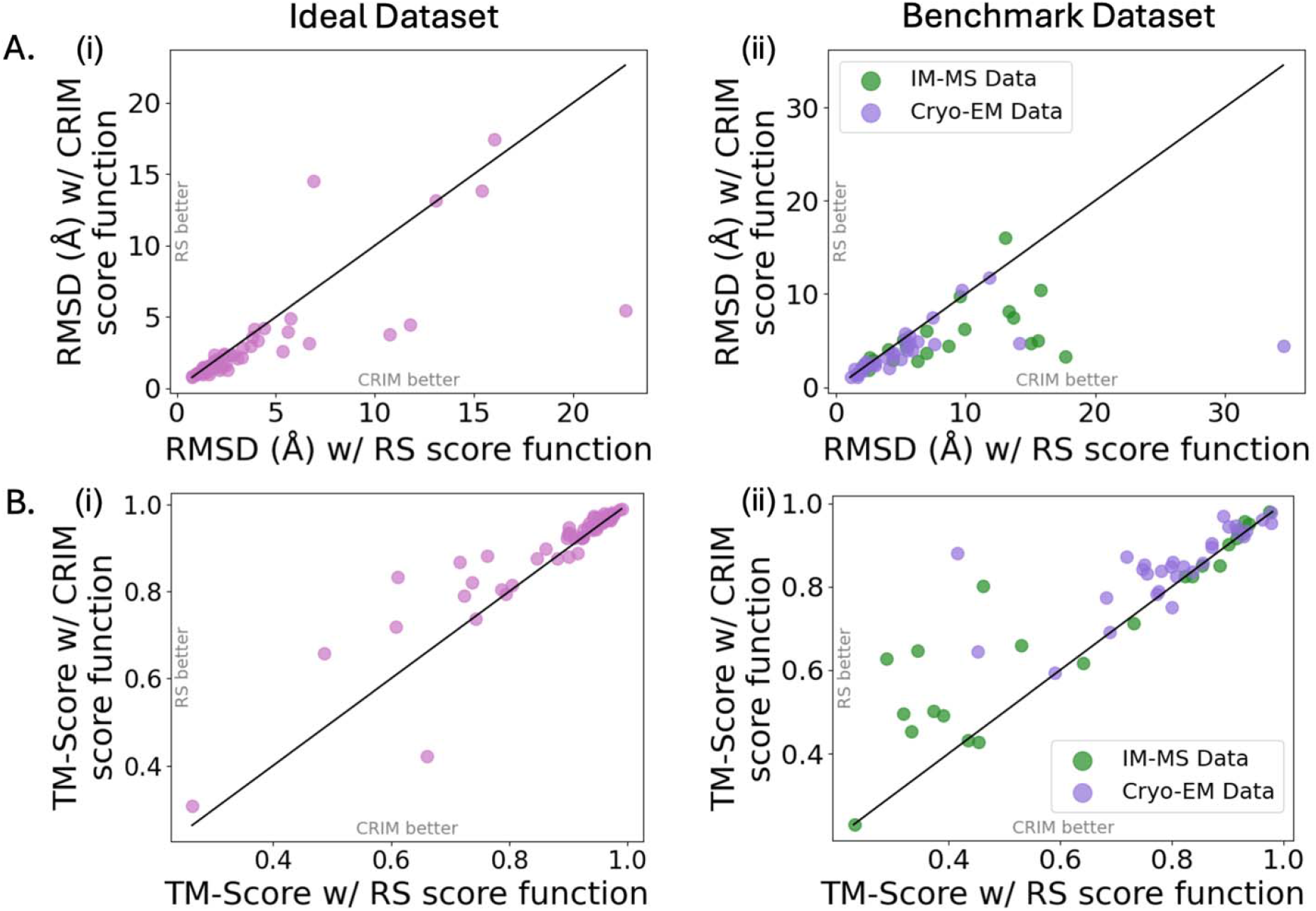
Evaluation of structural accuracy using CRIM versus RS score functions. (A) RMSD of the top-ranked models, presented for the ideal dataset (i) and the benchmark dataset (ii) separated by proteins with IM-MS data (green) and cryo-EM data (purple). (B) TM scores of the highest-ranking models, reported for the ideal dataset (i) and the benchmark dataset (ii) separated by proteins with IM-MS data (green) and cryo-EM data (purple).

### Cryo-EM and IM-MS Data Improved Model Selection in the Experimental Dataset

The experimental dataset consisted of 54 proteins with either experimental data from cryo-EM or IM-MS. The data was curated as described in the methods section. For each protein in the experimental dataset, 10,000 models were generated with Rosetta Comparative Modelling, and were rescored with the cryo-EM and IM-MS data.

Again, we saw an improvement in model selection quality with the addition of the CRIM score function as compared to the RS score function (Figure 2A (ii), 2B (ii)). The average best-model RMSD dropped from 6.65 Å (RS) to 4.38 Å (CRIM), and the mean TM-score rose from 0.73 (RS) to 0.79 (CRIM). Notably, most proteins benefited: 45/54 showed equal or lower RMSD with CRIM (Figure 2A (ii), and supplementary Fig. 4), and 43/54 showed equal or higher TM-scores (Figure 2B (ii), and supplementary Fig. 5). For the 45 proteins with lower RMSD values, we observed an average improvement of 3.40 Å (corresponding to an improvement of 0.09 in TM-score), with 20 of those achieving a greater than 1 Å RMSD reduction, and 17 achieving a greater than 0.05 TM-score increase, indicating widespread practical improvement with the CRIM score function. These results further demonstrate that model selection with cryo-EM and IM-MS data outperformed that of just the RS score function.

Furthermore, Figure 3 highlights two benchmark proteins, βB2-crystallin (PDBID: 1YTQ) and eEF3 (PDBID: 2IX8), where the CRIM score function significantly improved predictions over the RS score function. Figure 3A shows the RMSD vs score distributions of the models when scoring with the CRIM score function (orange) compared to scoring with the RS score function (blue). The corresponding TM-score distributions are shown in Figure 3B. We used an experimental CCS value for 1YTQ, whereas the cryo-EM density map was simulated as described in the methods section. 2IX8 had an experimental low-resolution cryo-EM density map, and the CCS value was simulated with PARCS. In both cases, we saw significantly better models selected when using the CRIM score function. Notably, both RS predictions had a TM-score below 0.5, which indicates that the predicted model did not share a similar fold to the native structure. However, when experimental data was included and the models were rescored with the CRIM score function, both predicted models exhibited a TM-score over 0.8, which indicates a highly similar structure between the predicted model and the native structure. This further supports that integrating these two types of experimental data does help in predicting the structures of monomeric proteins.

**Figure 3.**
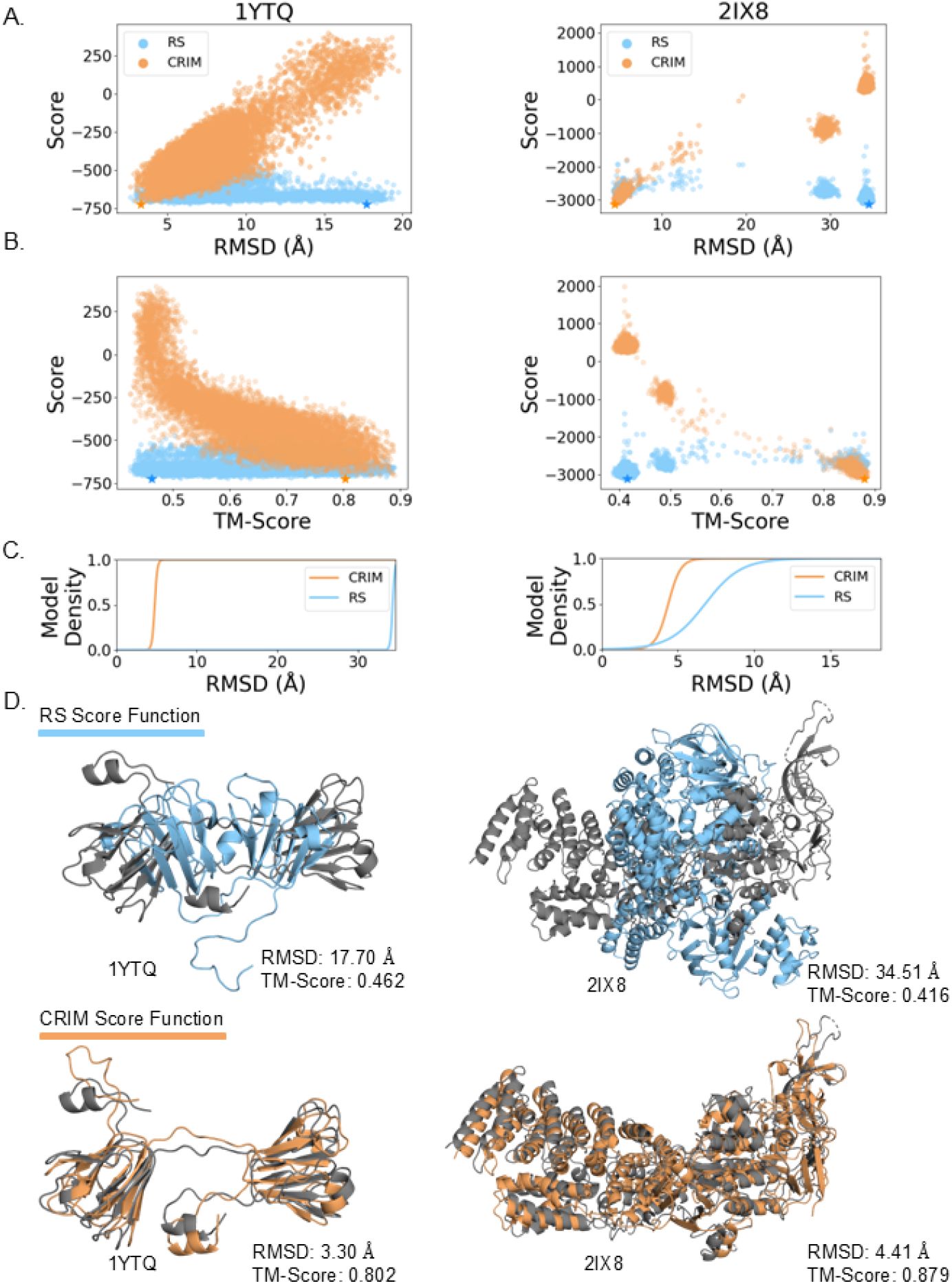
(A) Score versus RMSD distributions for the 10,000 model structures of 1YTQ and 2IX8 using the CRIM score function (orange) and the RS score function (light blue). For both score functions, the best scoring structure is indicated with a star in its respective color. (B) TM-score vs RMSD distributions of the models scored with the CRIM score function compared to the RS score function. (C) Cumulative density plots for the RMSD of top 100 scoring models. (D) The predicted models using the CRIM score function (orange) compared to the models predicted with the RS score function (light blue) with each of the models overlaid on the native structure (grey).

We performed a cumulative density plot analysis to understand if there was improvement in the 100 top scoring models’ RMSD as depicted in Figure 3C. For the two example proteins, the CRIM score function significantly shifted the distribution of the top 100 scoring models towards a lower RMSD value than when the RS score function was used. In both cases, the CRIM score function achieved a near complete density saturation at around 6 Å RMSD, whereas density saturation for models scored with the RS score function was only observed for RMSD values beyond 10 Å in both cases. This trend was seen throughout the experimental dataset (Supplementary Fig. 6), showing that the CRIM score function consistently scored models more accurately than the RS score function (even in cases where the top scoring model did not improve significantly).

### Structure Prediction Using CRIM is Competitive with AF3

AF3 is the state-of-the-art technique for protein structure prediction^15^. Thus, we compared the best CRIM predictions for the proteins in both the ideal and benchmark dataset to the respective AF3 predictions to contextualize the benefit of incorporating sparse experimental data into protein structure prediction (see Supplementary Fig. 7 for the full RMSD distributions across AF3, RS, and CRIM). For the ideal dataset (Figure 4A), the mean RMSD for the AF3 predictions was 2.55 Å, compared to a mean RMSD of 2.90 Å with the CRIM score function. While on average, AF3 slightly outperformed CRIM, 36/60 (60%) of the predictions were within 1.0 Å of each other, and for an additional 5 proteins CRIM outperformed AF3 by more than 1 Å RMSD. Thus, CRIM performed similarly or better than AF3 for 41/60 (68%) of the structures predicted as shown in Figure 4A. Similar trends were seen for the benchmark dataset comparison (Figure 4B). The mean RMSD for the AF3 predictions was 4.12 Å compared to 4.38 Å when using the CRIM score function. Across the dataset, 20/54 (37%) of the predictions were within 1 Å of each other, and for 12/54 proteins, CRIM produced lower RMSD models than AF3 by more than 1 Å RMSD. Notably, based on AF3’s reported training cutoff date of 09/30/2021, all proteins in the ideal dataset and all but two proteins in the benchmark dataset were potentially included in AF3 training; the two benchmark proteins deposited after this cutoff (8B2X and 7T7V) were both predicted substantially better by CRIM (RMSD values of 2.3 Å and 2.0 Å, respectively) than by AF3 (RMSD values of 19.3 Å and 5.0 Å, respectively). Collectively, these results demonstrate that although AF3 remains the more accurate method on average, CRIM frequently produced predictions that are competitive with AF3 and can substantially outperform AF3 for select targets when sparse experimental restraints provide strong discriminatory power. This observation is consistent with the broader motivation for CRIM. Experimentally grounded restraints reduce ambiguity and improve discrimination between native-like and non-native-like conformations, thereby improving predictions for difficult targets. Looking forward, an important extension of this work will be to incorporate sparse experimental information directly into machine learning based structure prediction workflows such as AF3, for example by using IM-MS and cryo-EM derived restraints during model generation and/or as part of a restraint-guided refinement stage. Integrating these orthogonal experimental constraints into machine learning pipelines could further reduce the frequency of inaccurate predictions and improve reliability for difficult targets, extending the benefits of sparse experimental data to next generation structure prediction workflows.

**Figure 4.**
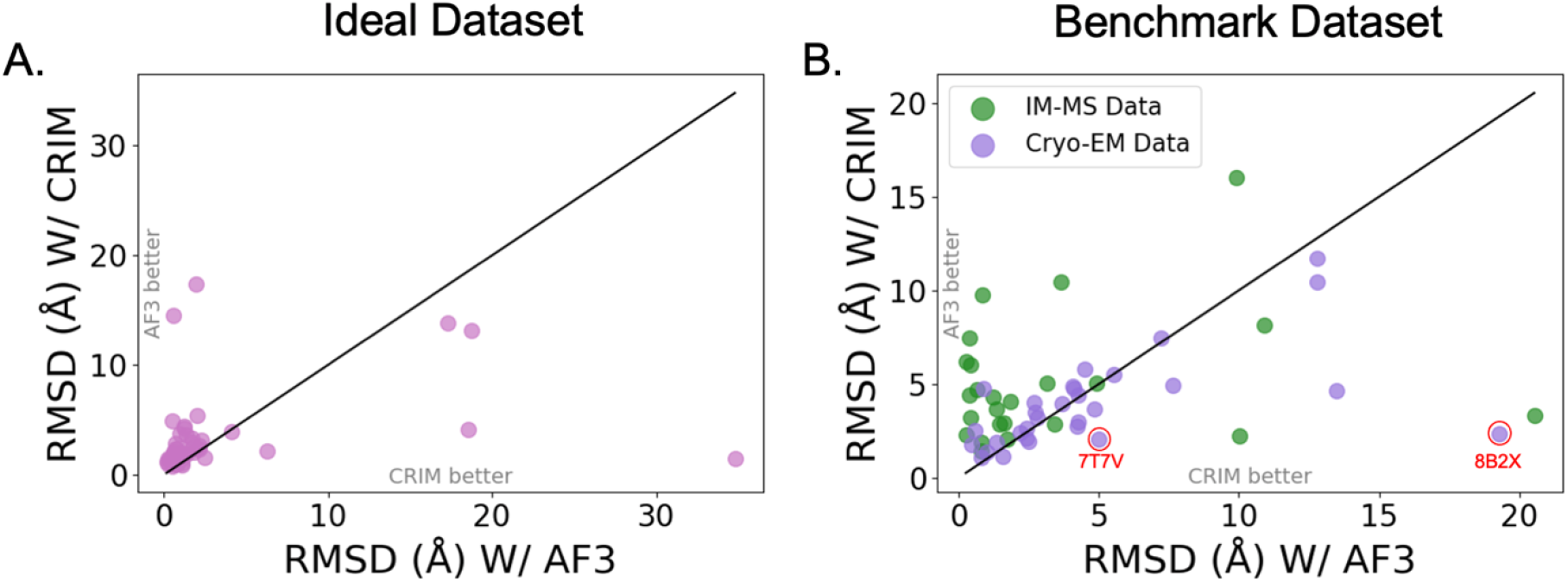
Comparison of structural accuracy when using the CRIM score function versus AF3. (A) RMSD of the top-ranked models for the ideal dataset. (B) RMSD of the top-ranked models for the benchmark dataset separated by proteins with IM-MS data (green) and cryo-EM data (purple). The two circled and labeled data points indicate the protein structures that were deposited in the PDB after the AF3 training date cutoff.

## Conclusion

This study introduced the Rosetta CRIM score function, an integrative scoring strategy that augments the RS score function with sparse cryo-EM density and IM-MS information to guide monomeric protein structure selection. Across an idealized, noise-free dataset of 60 proteins, CRIM consistently identified more native-like structures relative to RS alone, lowering the average best-model RMSD from 3.65 to 2.90 Å and increasing the mean TM-score from 0.88 to 0.90. In an experimental dataset of 54 proteins with either experimental cryo-EM or IM-MS data (and the complementary modality simulated), CRIM again improved quality: the mean best-model RMSD decreased from 6.65 to 4.38 Å and the mean TM-score rose from 0.73 to 0.79. The improvement was most pronounced for harder targets, where the additional information provided by IM-MS (shape information from CCS) and cryo-EM density helped to more clearly distinguish between native-like and non-native-like models. This clearly demonstrates the utility of integrative IM-MS/cryo-EM modeling for both ideal and experimental datasets. The CRIM score function is freely available as part of the Rosetta software suite. A detailed tutorial of the protein structure scoring pipeline is provided in the Supplementary information.

Moving forward, we plan to extend the CRIM score function beyond monomeric targets to explicitly handle protein complexes, where constraints from CCS and cryo-EM density can be integrated to improve discrimination among predicted architectures. Furthermore, incorporating CRIM into next-generation deep learning tools such as AlphaFold is an additional, natural extension of this work. AlphaFold frequently achieves high accuracy; however, it faces limitations for larger proteins, when evolutionary signals are weak, or for proteins with disordered regions and/or multiple conformations^55, 56^. In these settings, CRIM’s constraints from IM-MS data and low-resolution cryo-EM maps may add experimentally grounded information that helps reduce ambiguity and improves reliability for more difficult targets.

Finally, because CRIM combines IM-MS and electron microscopy information in a single framework, it may also be useful for modeling soft-landed proteins. Soft-landing workflows naturally pair native MS/IM-MS measurements with electron microscopy on the deposited sample^57-59^, which matches well with the types of data CRIM is designed to use. In this context, CRIM could help evaluate whether soft-landed proteins remain native-like and help select or refine structural models that agree with both the measured CCS value and the resulting EM density.

## Supporting information

Supplementary Figures 1-7 and tutorial

## Acknowledgements

We thank fellow Lindert lab members for their valuable discussions and guidance. We also acknowledge the computational resources provided by the Ohio Supercomputer Center (OSC)^60^. We are grateful to the Rosetta Commons community for developing and maintaining the Rosetta software suite and for making these tools broadly available. This work was supported by NIH (RM1-GM149374 to S.L.). We additionally acknowledge support from the NSF RaMP (Research and Mentoring for Postbacs; NSF 2216011 to S.L.) program for training, mentorship, and community resources that contributed to this work.

## References

(1) Coelho, E. D.; Arrais, J. P.; Oliveira, J. L. From protein-protein interactions to rational drug design: are computational methods up to the challenge? Curr Top Med Chem 2013, 13 (5), 602–618. DOI: 10.2174/1568026611313050005.

(2) Leelananda, S. P.; Lindert, S. Computational methods in drug discovery. Beilstein Journal of Organic Chemistry 2016, 12, 2694–2718. DOI: 10.3762/bjoc.12.267.

(3) Huang, P. S.; Boyken, S. E.; Baker, D. The coming of age of de novo protein design. Nature 2016, 537 (7620), 320–327. DOI: 10.1038/nature19946.

(4) Khoury, G. A.; Smadbeck, J.; Kieslich, C. A.; Floudas, C. A. Protein folding and de novo protein design for biotechnological applications. Trends Biotechnol 2014, 32 (2), 99–109. DOI: 10.1016/j.tibtech.2013.10.008.

(5) Thompson, M. C.; Yeates, T. O.; Rodriguez, J. A. Advances in methods for atomic resolution macromolecular structure determination. F1000Res 2020, 9. DOI: 10.12688/f1000research.25097.1.

(6) Robertson, J. C.; Nassar, R.; Liu, C.; Brini, E.; Dill, K. A.; Perez, A. NMR-assisted protein structure prediction with MELDxMD. Proteins 2019, 87 (12), 1333–1340. DOI: 10.1002/prot.25788.

(7) Weiner, B. E.; Alexander, N.; Akin, L. R.; Woetzel, N.; Karakas, M.; Meiler, J. BCL::Fold--protein topology determination from limited NMR restraints. Proteins 2014, 82 (4), 587–595. DOI: 10.1002/prot.24427.

(8) Vernon, R.; Shen, Y.; Baker, D.; Lange, O. F. Improved chemical shift based fragment selection for CS-Rosetta using Rosetta3 fragment picker. J Biomol NMR 2013, 57 (2), 117–127. DOI: 10.1007/s10858-013-9772-4.

(9) Frenz, B.; Walls, A. C.; Egelman, E. H.; Veesler, D.; DiMaio, F. RosettaES: a sampling strategy enabling automated interpretation of difficult cryo-EM maps. Nat Methods 2017, 14 (8), 797–800. DOI: 10.1038/nmeth.4340.

(10) Lindert, S.; Alexander, N.; Wötzel, N.; Karakaş, M.; Stewart, P. L.; Meiler, J. EM-fold: de novo atomic-detail protein structure determination from medium-resolution density maps. Structure 2012, 20 (3), 464–478. DOI: 10.1016/j.str.2012.01.023.

(11) Artigues, A.; Nadeau, O. W.; Rimmer, M. A.; Villar, M. T.; Du, X.; Fenton, A. W.; Carlson, G. M. Protein Structural Analysis via Mass Spectrometry-Based Proteomics. Adv Exp Med Biol 2016, 919, 397–431. DOI: 10.1007/978-3-319-41448-5_19.

(12) Stiving, A. Q.; VanAernum, Z. L.; Busch, F.; Harvey, S. R.; Sarni, S. H.; Wysocki, V. H. Surface-Induced Dissociation: An Effective Method for Characterization of Protein Quaternary Structure. Anal Chem 2019, 91 (1), 190–209. DOI: 10.1021/acs.analchem.8b05071.

(13) Konijnenberg, A.; Butterer, A.; Sobott, F. Native ion mobility-mass spectrometry and related methods in structural biology. Biochim Biophys Acta 2013, 1834 (6), 1239–1256. DOI: 10.1016/j.bbapap.2012.11.013.

(14) Marklund, E. G.; Degiacomi, M. T.; Robinson, C. V.; Baldwin, A. J.; Benesch, J. L. Collision cross sections for structural proteomics. Structure 2015, 23 (4), 791–799. DOI: 10.1016/j.str.2015.02.010.

(15) Abramson, J.; Adler, J.; Dunger, J.; Evans, R.; Green, T.; Pritzel, A.; Ronneberger, O.; Willmore, L.; Ballard, A. J.; Bambrick, J.; et al. Accurate structure prediction of biomolecular interactions with AlphaFold 3. Nature 2024, 630 (8016), 493–500. DOI: 10.1038/s41586-024-07487-w.

(16) Wohlwend, J.; Corso, G.; Passaro, S.; Reveiz, M.; Leidal, K.; Swiderski, W.; Portnoi, T.; Chinn, I.; Silterra, J.; Jaakkola, T.; et al. Boltz-1 Democratizing Biomolecular Interaction Modeling. bioRxiv 2024. DOI: 10.1101/2024.11.19.624167.

(17) Baek, M.; DiMaio, F.; Anishchenko, I.; Dauparas, J.; Ovchinnikov, S.; Lee, G. R.; Wang, J.; Cong, Q.; Kinch, L. N.; Schaeffer, R. D.; et al. Accurate prediction of protein structures and interactions using a three-track neural network. Science 2021, 373 (6557), 871–876. DOI: 10.1126/science.abj8754.

(18) Agarwal, V.; McShan, A. C. The power and pitfalls of AlphaFold2 for structure prediction beyond rigid globular proteins. Nat Chem Biol 2024, 20 (8), 950–959. DOI: 10.1038/s41589-024-01638-w.

(19) Seffernick, J. T.; Lindert, S. Hybrid methods for combined experimental and computational determination of protein structure. J Chem Phys 2020, 153 (24), 240901. DOI: 10.1063/5.0026025.

(20) Biehn, S. E.; Lindert, S. Protein Structure Prediction with Mass Spectrometry Data. Annu Rev Phys Chem 2022, 73, 1–19. DOI: 10.1146/annurev-physchem-082720-123928 From NLM.

(21) Leman, J. K.; Weitzner, B. D.; Lewis, S. M.; Adolf-Bryfogle, J.; Alam, N.; Alford, R. F.; Aprahamian, M.; Baker, D.; Barlow, K. A.; Barth, P.; et al. Macromolecular modeling and design in Rosetta: recent methods and frameworks. Nat Methods 2020, 17 (7), 665–680. DOI: 10.1038/s41592-020-0848-2.

(22) Bonvin, P. I. K. a. A. M. J. J. Integrative Modelling of Biomolecular Complexes. Journal of Molecular Biology 2020, 432 (9), 2861–2881. DOI: 10.1016/j.jmb.2019.11.009.

(23) Bullock, J. M. A.; Sen, N.; Thalassinos, K.; Topf, M. Modeling Protein Complexes Using Restraints from Crosslinking Mass Spectrometry. Structure 2018, 26 (7), 1015–1024.e1012. DOI: 10.1016/j.str.2018.04.016.

(24) Khakzad, H.; Vermeul, S.; Malmström, L. Quaternary Structure Modeling Through Chemical Cross-Linking Mass Spectrometry: Extending TX-MS Jupyter Reports. J Vis Exp 2021, (176). DOI: 10.3791/60311.

(25) Aprahamian, M. L.; Chea, E. E.; Jones, L. M.; Lindert, S. Rosetta Protein Structure Prediction from Hydroxyl Radical Protein Footprinting Mass Spectrometry Data. Anal Chem 2018, 90 (12), 7721–7729. DOI: 10.1021/acs.analchem.8b01624.

(26) Aprahamian, M. L.; Lindert, S. Utility of Covalent Labeling Mass Spectrometry Data in Protein Structure Prediction with Rosetta. J Chem Theory Comput 2019, 15 (5), 3410–3424. DOI: 10.1021/acs.jctc.9b00101.

(27) Day, E. H.; Lindert, S. Extracting Residue Solvent Exposure from Covalent Labeling Data with Machine Learning: A Hybrid Approach for Protein Structure Prediction. J Am Soc Mass Spectrom 2025, 36 (6), 1336–1347. DOI: 10.1021/jasms.5c00053.

(28) Biehn, S. E.; Limpikirati, P.; Vachet, R. W.; Lindert, S. Utilization of Hydrophobic Microenvironment Sensitivity in Diethylpyrocarbonate Labeling for Protein Structure Prediction. Anal Chem 2021, 93 (23), 8188–8195. DOI: 10.1021/acs.analchem.1c00395.

(29) Schmidt, C.; Macpherson, J. A.; Lau, A. M.; Tan, K. W.; Fraternali, F.; Politis, A. Surface Accessibility and Dynamics of Macromolecular Assemblies Probed by Covalent Labeling Mass Spectrometry and Integrative Modeling. Anal Chem 2017, 89 (3), 1459–1468. DOI: 10.1021/acs.analchem.6b02875.

(30) Bolz, R. M.; Seffernick, J. T.; Drake, Z. C.; Harvey, S. R.; Wysocki, V. H.; Lindert, S. Energy Resolved Mass Spectrometry Data from Surfaced Induced Dissociation Improves Prediction of Protein Complex Structure. Anal Chem 2025, 97 (4), 2375–2383. DOI: 10.1021/acs.analchem.4c05837.

(31) Seffernick, J. T.; Harvey, S. R.; Wysocki, V. H.; Lindert, S. Predicting Protein Complex Structure from Surface-Induced Dissociation Mass Spectrometry Data. ACS Cent Sci 2019, 5 (8), 1330–1341. DOI: 10.1021/acscentsci.8b00912.

(32) Seffernick, J. T.; Canfield, S. M.; Harvey, S. R.; Wysocki, V. H.; Lindert, S. Prediction of Protein Complex Structure Using Surface-Induced Dissociation and Cryo-Electron Microscopy. Anal Chem 2021, 93 (21), 7596–7605. DOI: 10.1021/acs.analchem.0c05468.

(33) Snyder, D. T.; Harvey, S. R.; Wysocki, V. H. Surface-induced Dissociation Mass Spectrometry as a Structural Biology Tool. Chem Rev 2022, 122 (8), 7442–7487. DOI: 10.1021/acs.chemrev.1c00309.

(34) Sahasrabuddhe, A.; Hsia, Y.; Busch, F.; Sheffler, W.; King, N. P.; Baker, D.; Wysocki, V. H. Confirmation of intersubunit connectivity and topology of designed protein complexes by native MS. Proc Natl Acad Sci U S A 2018, 115 (6), 1268–1273. DOI: 10.1073/pnas.1713646115.

(35) Degiacomi, M. T. On the Effect of Sphere-Overlap on Super Coarse-Grained Models of Protein Assemblies. J Am Soc Mass Spectrom 2019, 30 (1), 113–117. DOI: 10.1007/s13361-018-1974-2.

(36) Eschweiler, J. D.; Frank, A. T.; Ruotolo, B. T. Coming to Grips with Ambiguity: Ion Mobility-Mass Spectrometry for Protein Quaternary Structure Assignment. J Am Soc Mass Spectrom 2017, 28 (10), 1991–2000. DOI: 10.1007/s13361-017-1757-1.

(37) Jurneczko, E.; Barran, P. E. How useful is ion mobility mass spectrometry for structural biology? The relationship between protein crystal structures and their collision cross sections in the gas phase. Analyst 2011, 136 (1), 20–28. DOI: 10.1039/c0an00373e.

(38) McCabe, J. W.; Mallis, C. S.; Kocurek, K. I.; Poltash, M. L.; Shirzadeh, M.; Hebert, M. J.; Fan, L.; Walker, T. E.; Zheng, X.; Jiang, T.; et al. First-Principles Collision Cross Section Measurements of Large Proteins and Protein Complexes. Anal Chem 2020, 92 (16), 11155–11163. DOI: 10.1021/acs.analchem.0c01285.

(39) Politis, A.; Park, A. Y.; Hall, Z.; Ruotolo, B. T.; Robinson, C. V. Integrative modelling coupled with ion mobility mass spectrometry reveals structural features of the clamp loader in complex with single-stranded DNA binding protein. J Mol Biol 2013, 425 (23), 4790–4801. DOI: 10.1016/j.jmb.2013.04.006.

(40) Turzo, S. M. B. A.; Seffernick, J. T.; Rolland, A. D.; Donor, M. T.; Heinze, S.; Prell, J. S.; Wysocki, V. H.; Lindert, S. Protein shape sampled by ion mobility mass spectrometry consistently improves protein structure prediction. Nat Commun 2022, 13 (1), 4377. DOI: 10.1038/s41467-022-32075-9.

(41) Eschweiler, J. D.; Farrugia, M. A.; Dixit, S. M.; Hausinger, R. P.; Ruotolo, B. T. A Structural Model of the Urease Activation Complex Derived from Ion Mobility-Mass Spectrometry and Integrative Modeling. Structure 2018, 26 (4), 599–606.e593. DOI: 10.1016/j.str.2018.03.001.

(42) Lindert, S.; Staritzbichler, R.; Wötzel, N.; Karakaş, M.; Stewart, P. L.; Meiler, J. EM-fold: De novo folding of alpha-helical proteins guided by intermediate-resolution electron microscopy density maps. Structure 2009, 17 (7), 990–1003. DOI: 10.1016/j.str.2009.06.001.

(43) Lindert, S.; Hofmann, T.; Wötzel, N.; Karakaş, M.; Stewart, P. L.; Meiler, J. Ab initio protein modeling into CryoEM density maps using EM-Fold. Biopolymers 2012, 97 (9), 669–677. DOI: 10.1002/bip.22027.

(44) Wang, R. Y.; Song, Y.; Barad, B. A.; Cheng, Y.; Fraser, J. S.; DiMaio, F. Automated structure refinement of macromolecular assemblies from cryo-EM maps using Rosetta. Elife 2016, 5. DOI: 10.7554/eLife.17219.

(45) Lau, A. M.; Politis, A. Integrative Mass Spectrometry-Based Approaches for Modeling Macromolecular Assemblies. Methods Mol Biol 2021, 2247, 221–241. DOI: 10.1007/978-1-0716-1126-5_12.

(46) Sali, A. From integrative structural biology to cell biology. Journal of Biological Chemistry 2021, 296. DOI: 10.1016/j.jbc.2021.100743 (accessed 2025/10/28).

(47) Turzo, S. M. B. A.; Seffernick, J. T.; Lyskov, S.; Lindert, S. Predicting ion mobility collision cross sections using projection approximation with ROSIE-PARCS webserver. Brief Bioinform 2023, 24 (5). DOI: 10.1093/bib/bbad308.

(48) Song, Y.; DiMaio, F.; Wang, R. Y.; Kim, D.; Miles, C.; Brunette, T.; Thompson, J.; Baker, D. High-resolution comparative modeling with RosettaCM. Structure 2013, 21 (10), 1735–1742. DOI: 10.1016/j.str.2013.08.005 From NLM Medline.

(49) Gront, D.; Kulp, D. W.; Vernon, R. M.; Strauss, C. E.; Baker, D. Generalized fragment picking in Rosetta: design, protocols and applications. PLoS One 2011, 6 (8), e23294. DOI: 10.1371/journal.pone.0023294 From NLM Medline.

(50) Alford, R. F.; Leaver-Fay, A.; Jeliazkov, J. R.; O’Meara, M. J.; DiMaio, F. P.; Park, H.; Shapovalov, M. V.; Renfrew, P. D.; Mulligan, V. K.; Kappel, K.; et al. The Rosetta All-Atom Energy Function for Macromolecular Modeling and Design. J Chem Theory Comput 2017, 13 (6), 3031–3048. DOI: 10.1021/acs.jctc.7b00125 From NLM Medline.

(51) Zhang, Y.; Skolnick, J. TM-align: a protein structure alignment algorithm based on the TM-score. Nucleic Acids Res 2005, 33 (7), 2302–2309. DOI: 10.1093/nar/gki524 From NLM Medline.

(52) Meng, E. C.; Goddard, T. D.; Pettersen, E. F.; Couch, G. S.; Pearson, Z. J.; Morris, J. H.; Ferrin, T. E. UCSF ChimeraX: Tools for structure building and analysis. Protein Sci 2023, 32 (11), e4792. DOI: 10.1002/pro.4792.

(53) Woetzel, N.; Lindert, S.; Stewart, P. L.; Meiler, J. BCL::EM-Fit: rigid body fitting of atomic structures into density maps using geometric hashing and real space refinement. J Struct Biol 2011, 175 (3), 264–276. DOI: 10.1016/j.jsb.2011.04.016.

(54) DiMaio, F.; Tyka, M. D.; Baker, M. L.; Chiu, W.; Baker, D. Refinement of protein structures into low-resolution density maps using rosetta. J Mol Biol 2009, 392 (1), 181–190. DOI: 10.1016/j.jmb.2009.07.008 From NLM Medline.

(55) Saldaño, T.; Escobedo, N.; Marchetti, J.; Zea, D. J.; Mac Donagh, J.; Velez Rueda, A. J.; Gonik, E.; García Melani, A.; Novomisky Nechcoff, J.; Salas, M. N.; et al. Impact of protein conformational diversity on AlphaFold predictions. Bioinformatics 2022, 38 (10), 2742–2748. DOI: 10.1093/bioinformatics/btac202.

(56) Chakravarty, D.; Schafer, J. W.; Chen, E. A.; Thole, J. F.; Ronish, L. A.; Lee, M.; Porter, L. L. AlphaFold predictions of fold-switched conformations are driven by structure memorization. Nat Commun 2024, 15 (1), 7296. DOI: 10.1038/s41467-024-51801-z.

(57) Esser, T. K.; Böhning, J.; Fremdling, P.; Bharat, T.; Gault, J.; Rauschenbach, S. Cryo-EM samples of gas-phase purified protein assemblies using native electrospray ion-beam deposition. Faraday Discuss 2022, 240 (0), 67–80. DOI: 10.1039/d2fd00065b.

(58) Esser, T. K.; Böhning, J.; Önür, A.; Chinthapalli, D. K.; Eriksson, L.; Grabarics, M.; Fremdling, P.; Konijnenberg, A.; Makarov, A.; Botman, A.; et al. Cryo-EM of soft-landed β-galactosidase: Gas-phase and native structures are remarkably similar. Sci Adv 2024, 10 (7), eadl4628. DOI: 10.1126/sciadv.adl4628.

(59) Westphall, M. S.; Lee, K. W.; Hemme, C.; Salome, A. Z.; Mertz, K.; Grant, T.; Coon, J. J. Cryogenic Soft Landing Improves Structural Preservation of Protein Complexes. Anal Chem 2023, 95 (40), 15094–15101. DOI: 10.1021/acs.analchem.3c03228.

(60) Ohio Supercomputer, C. Ohio Supercomputer Center. 1987.

