## Supplementary Figures 1-7 and tutorial for "Improving Protein Structure Prediction Using Integrative Cryo-EM and Ion Mobility Mass Spectrometry Modeling"

Supplementary Figure 1………………..……………………………………………………….....2

Supplementary Figure 2………………..……………………………………………………….....4

Supplementary Figure 3………………..……………………………………………………….....6

Supplementary Figure 4………………..……………………………………………………….....8

Supplementary Figure 5………………..……………………………………………....………...10

Supplementary Figure 6………………..………………………………………………......…….12

Supplementary Figure 7………………..………………………………………………………...13

Tutorial...........................................................................................................................................14

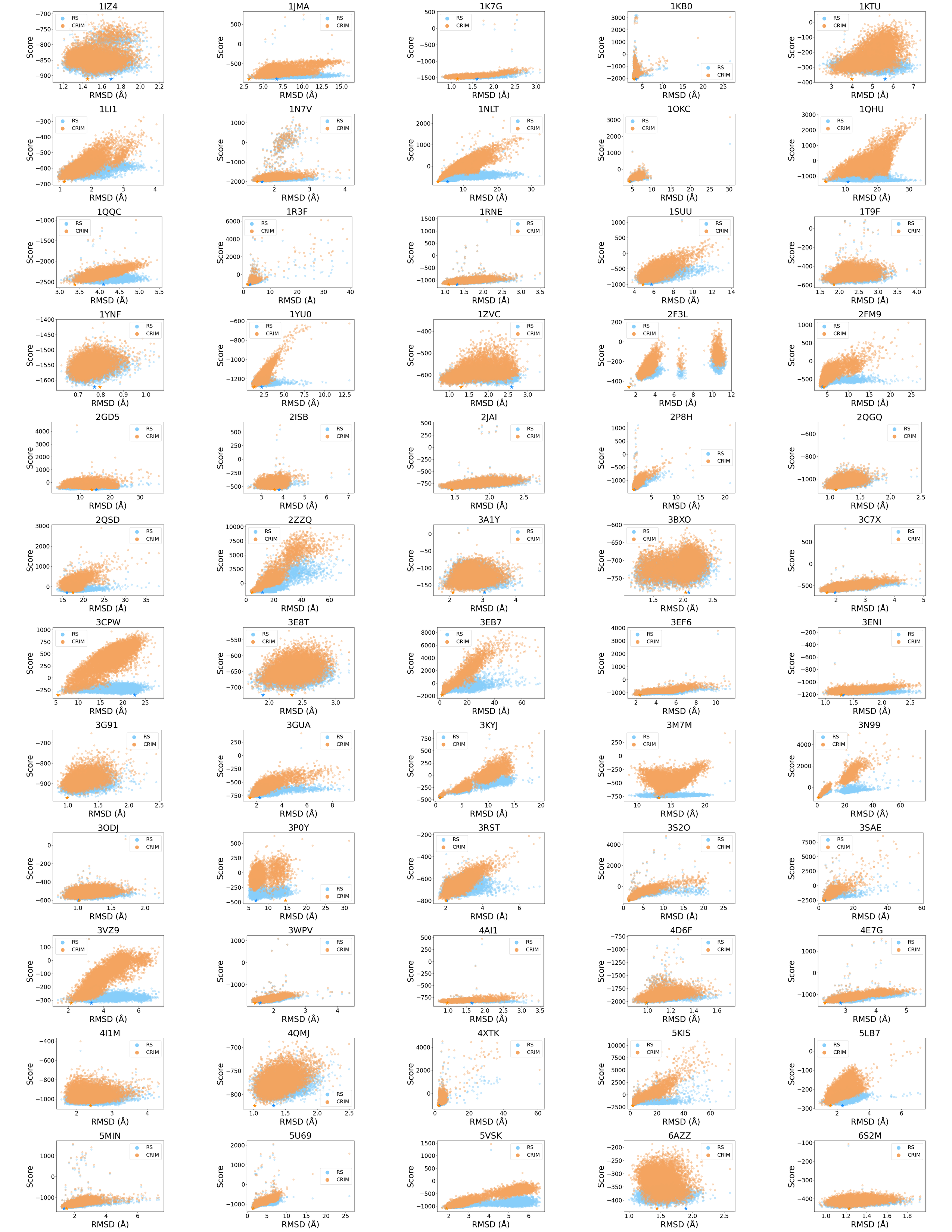

Supplementary Figure 1. Score versus RMSD for the 10,000 model structures for each of the ideal dataset proteins using the RS score function (light blue) and the CRIM score function (orange). For both score functions, the best scoring structure is indicated with a star in its respective color. The score values for the CRIM score function were normalized to the RS scores by subtracting the difference between the two best scoring structures from all 10,000 of the model scores.

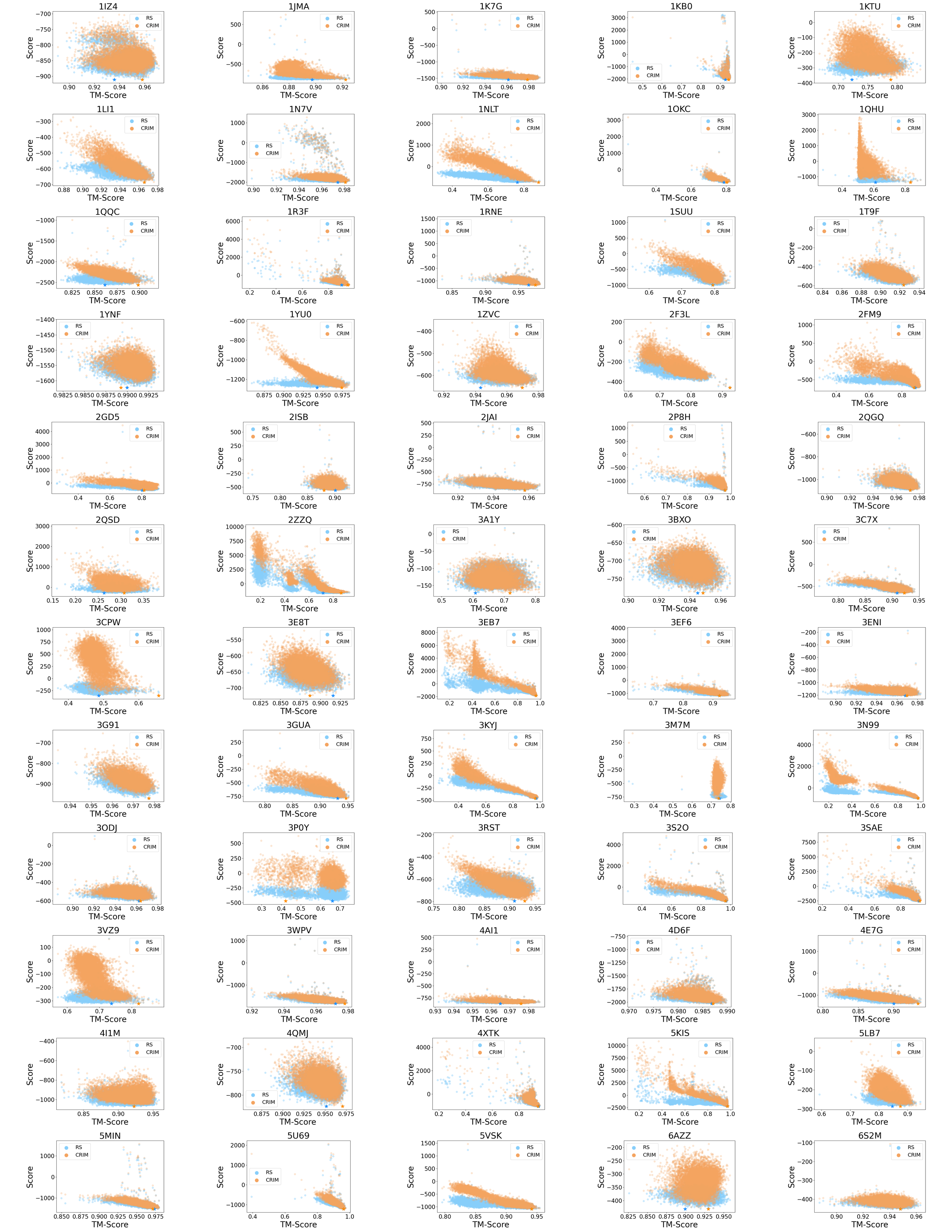

Supplementary Figure 2. Score versus TM-score for the 10,000 model structures for each of the ideal dataset proteins using the RS score function (light blue) and the CRIM score function (orange). For both score functions, the best scoring structure is indicated with a star in its respective color. The score values for the CRIM score function were normalized to the RS scores by subtracting the difference between the two best scoring structures from all 10,000 of the model scores.

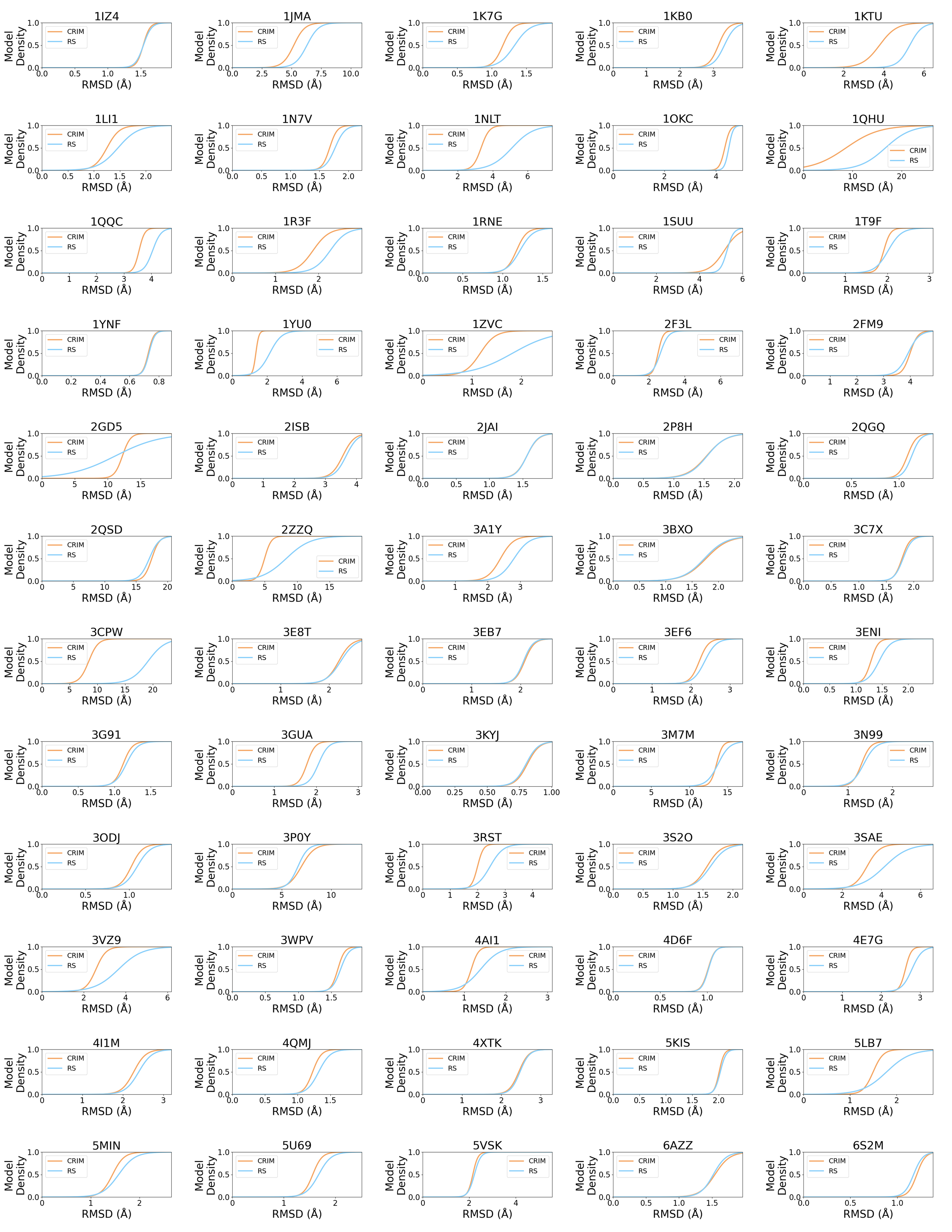

Supplementary Figure 3. Cumulative density plots for the RMSD of the top 100 scoring models in the ideal dataset. The structures scored with the RS score function are depicted in light blue, while the structures scored with CRIM are depicted in orange.

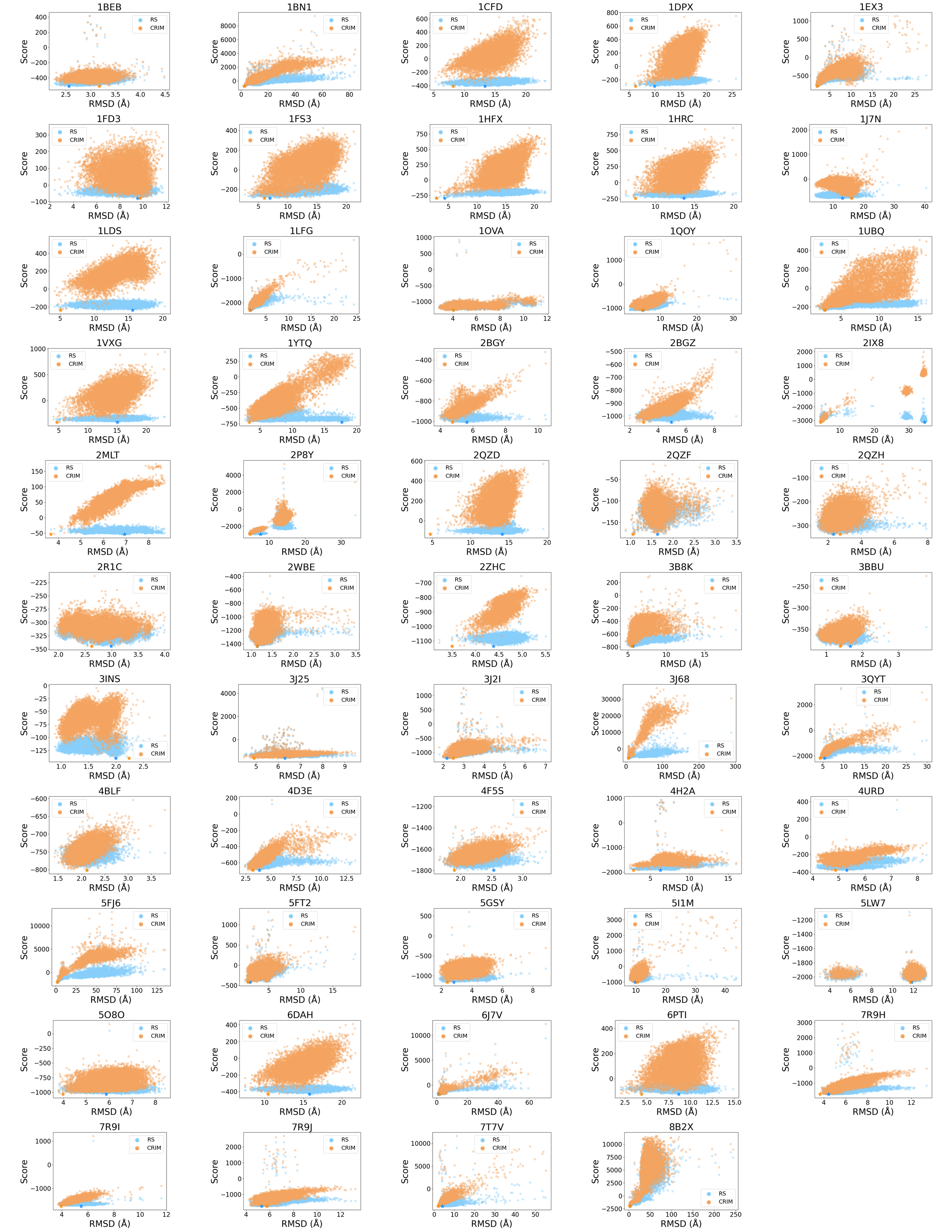

Supplementary Figure 4. Score versus RMSD for the 10,000 model structures for each of the benchmark dataset proteins using the RS score function (light blue) and the CRIM score function (orange). For both score functions, the best scoring structure is indicated with a star in its respective color. The score values for the CRIM score function were normalized to the RS scores by subtracting the difference between the two best scoring structures from all 10,000 of the model scores.

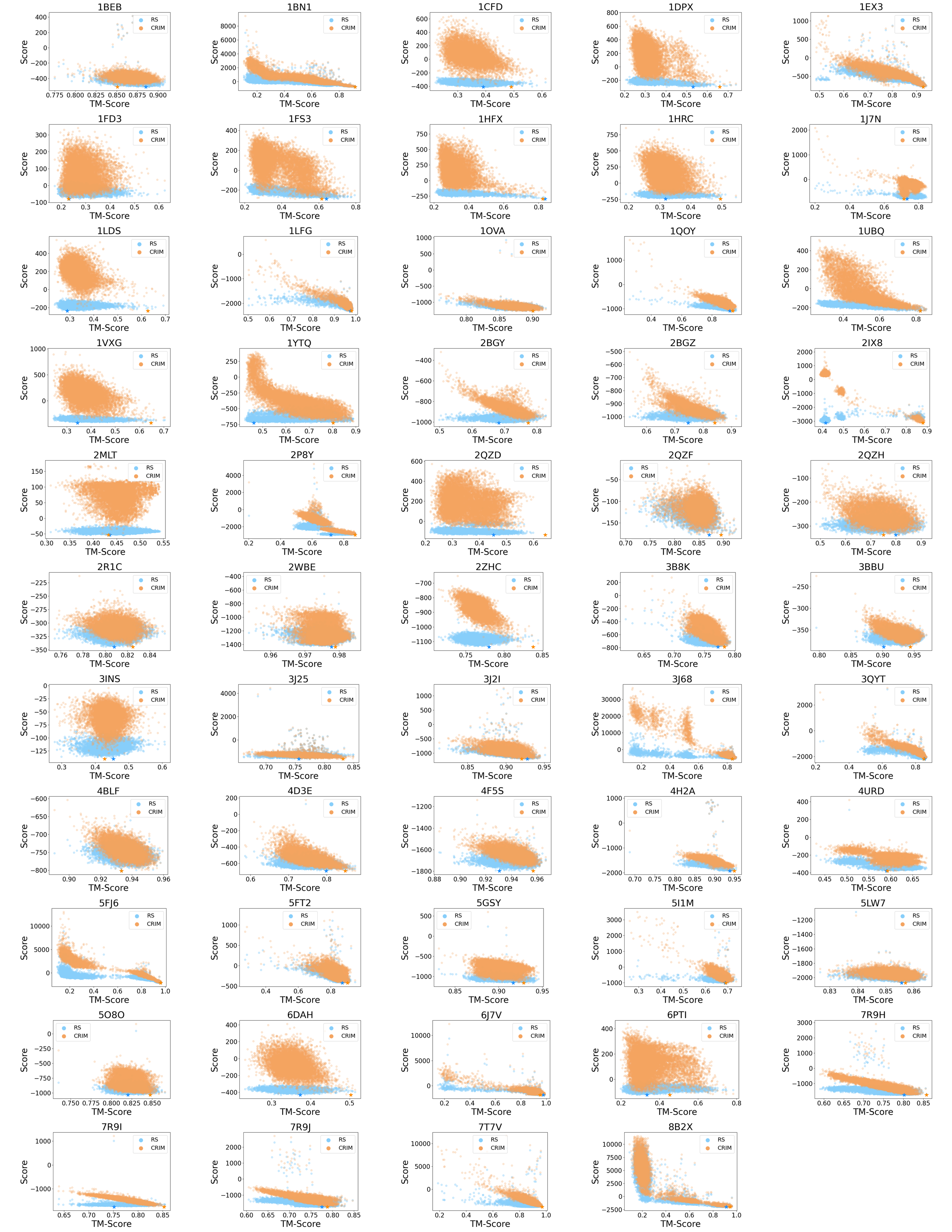

Supplementary Figure 5. Score versus TM-scores for the 10,000 model structures for each of the benchmark dataset proteins using the RS score function (light blue) and the CRIM score function (orange). For both score functions, the best scoring structure is indicated with a star in its respective color. The score values for the CRIM score function were normalized to the RS scores by subtracting the difference between the two best scoring structures from all 10,000 of the model scores.

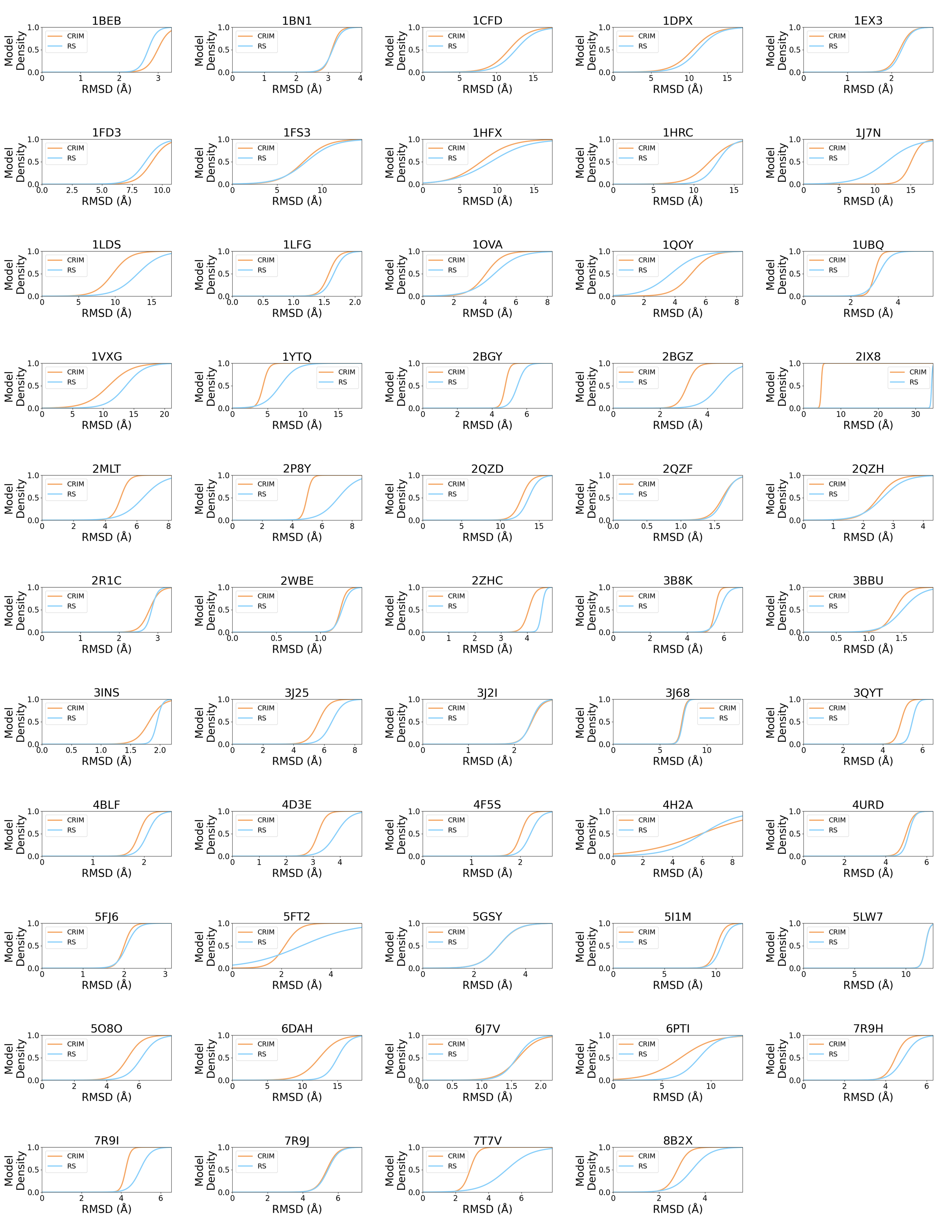

Supplementary Figure 6. Cumulative density plots for the RMSD of the top 100 scoring models in the benchmark dataset. The structures scored with the RS score function are depicted in light blue, while the structures scored with CRIM are depicted in orange.

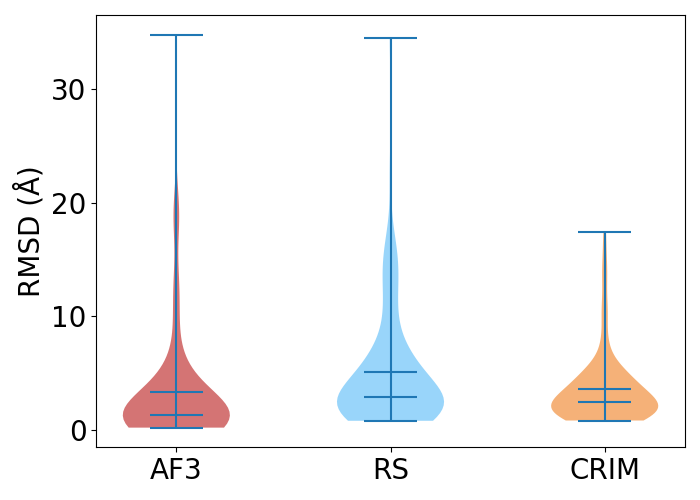

Supplementary Figure 7. Comparison of the RMSDs of the best scoring models identified with different score terms to the AlphaFold3 predictions for all proteins across both the experimental and ideal datasets (red). The RS predictions are the structures predicted with RosettaCM, scored with REF2015 score (light blue). The CRIM predictions represent predictions that were scored with the CRIM score (orange).

### **Tutorial**

### **1. Density Map Generation (BCL)**

Cryo-EM density maps were simulated from PDB structures using the BioChemical Library (BCL). These maps are later used for Rosetta cryo-EM scoring.^1, 2^

~/bcl-3.6.1-Source/build/linux64_release/bin/bcl-apps-static.exe \

density:FromPDB pdb_file.pdb \

-resolution 14.0 \

-voxel_size 1.4 \

-kernel GaussianSphere \

-noise 0.8 \

-random_seed

Here,

- pdb_file.pdb is the atomic model of the protein.
- The voxel size is typically set to **1/10 of the resolution**.
- Resolution, voxel size, and noise level can be adjusted depending on the system.
- Alternatively, experimental cryo-EM maps can be used directly.

### **2. Cryo-EM Density Scoring in Rosetta**

Cryo-EM map density agreement was evaluated using Rosetta’s electron density scoring framework, as implemented in the Rosetta software suite (<https://github.com/RosettaCommons/rosetta/>).^3^

~Rosetta/main/source/bin/score_jd2.default.linuxgccrelease \

@flags_cryoEM_score

-database ~/Rosetta/main/database/

-in:file:s relaxed_pdb_file.pdb

-ex1

-ex2aro

-edensity::mapfile density_map_name.mrc

-edensity::mapreso 14.0

-edensity::cryoem_scatterers

-crystal_refine

-restore_talaris_behavior

-out:prefix cryoEM_score

In this framework, agreement between the model and the cryo-EM map is reported via the elec_dens_fast term in the Rosetta score file. Density maps used for scoring can either be simulated from atomic models or obtained directly from experimental cryo-EM data. This score term is referred to as the ${"Cryo-EM}_{Score Term}"$ in the ${CRIM}_{Score}$ function.

**3. IM-MS scoring in Rosetta**

<path/to/Rosetta>/main/source/bin/score.default.<os><compiler>release \

-database <path/to/Rosetta>/main/database \

-in:file:s model.pdb \

-ccs_nrots 300 \

-ccs_prad 1.0 \

-ccs_exp experimental_ccs_value \

-score:patch ccs_imms_cryoem.wts_patch

Here, the experimentally measured CCS value is provided via the -ccs_exp flag. Upon successful execution, the weighted IM-MS contribution is reported in the “ccs_imms_cryoem” column of the Rosetta score file. The “score” column corresponds to the standard REF2015 energy function^4^ with the IM _Score Term_ ($RS+2\left( {IM}_{Score Term} \right)$) added.

### **4. Combined IM-MS and Cryo-EM Scoring**

To jointly evaluate structural models against both IM-MS and cryo-EM data, the combined scoring function, ${CRIM}_{Score}$, was employed.

This integrated approach leverages complementary experimental information by combining the individual scoring terms obtained from cryo-EM density fitting and IM-MS CCS agreement.

The scores derived from cryo-EM density scoring (Section 2) and IM-MS scoring (Section 3) were linearly combined to compute the CRIM score as described in the equation 2 in the manuscript. The final combined score was calculated as a weighted sum of the individual terms, using predefined weights to balance the relative contributions of cryo-EM and IM-MS information.

References

(1) Meng, E. C.; Goddard, T. D.; Pettersen, E. F.; Couch, G. S.; Pearson, Z. J.; Morris, J. H.; Ferrin, T. E. UCSF ChimeraX: Tools for structure building and analysis. *Protein Sci* **2023**, *32* (11), e4792. DOI: 10.1002/pro.4792.

(2) Woetzel, N.; Lindert, S.; Stewart, P. L.; Meiler, J. BCL::EM-Fit: rigid body fitting of atomic structures into density maps using geometric hashing and real space refinement. *J Struct Biol* **2011**, *175* (3), 264–276. DOI: 10.1016/j.jsb.2011.04.016.

(3) DiMaio, F.; Tyka, M. D.; Baker, M. L.; Chiu, W.; Baker, D. Refinement of protein structures into low-resolution density maps using rosetta. *J Mol Biol* **2009**, *392* (1), 181–190. DOI: 10.1016/j.jmb.2009.07.008 From NLM Medline.

(4) Alford, R. F.; Leaver-Fay, A.; Jeliazkov, J. R.; O'Meara, M. J.; DiMaio, F. P.; Park, H.; Shapovalov, M. V.; Renfrew, P. D.; Mulligan, V. K.; Kappel, K.; et al. The Rosetta All-Atom Energy Function for Macromolecular Modeling and Design. *J Chem Theory Comput* **2017**, *13*
